# Overnight Th1 immune activation predicts the cortisol awakening response

**DOI:** 10.64898/2026.07.06.735751

**Authors:** Lennart Seizer, Michaela Garlík Matušková, Johanna M. Gostner, Christian Schubert

**Affiliations:** Institute of General Practice and Family Medicine, LMU University Hospital, Germany; Department of Child and Adolescent Psychiatry, Psychosomatics and Psychotherapy, University Hospital of Tübingen, Germany; German Center for Mental Health (DZPG), Germany; Institute of Medical Biochemistry, Biocenter, Medical University Innsbruck, Austria; Department of Pharmaceutical Analysis and Nuclear Pharmacy, Faculty of Pharmacy, Comenius University Bratislava, Slovakia; Department of Clinical Pharmacology, Faculty Hospital in Trnava, Slovakia; Department of Psychiatry, Psychotherapy, Psychosomatics and Medical Psychology, Medical University Innsbruck, Austria

**Keywords:** cortisol awakening response, neopterin, diurnal, circadian, immune system, time series, ambulatory assessment, intensive longitudinal data, psychoneuroimmunology

## Abstract

The cortisol awakening response (CAR) marks the transition from rest to wake phase by a sharp increase in cortisol levels upon awakening in the morning. This increase may assist in cognitive and behavioral awakening, but its function is not fully understood yet. In this pilot study we aimed to provide first data on the influence of immune system activity on the CAR. Thereby, a within-subject analysis approach was applied to avoid confounding by between-subject bias and improve inference of the results. Three healthy subjects collected their overnight urine for analysis of neopterin (Th1 immune activation marker) and sampled saliva at 0, 30, and 45 minutes after awakening in the morning for cortisol determination and CAR estimation. Additionally, subjects wore an EEG-headband overnight for objective determination of the awakening timepoint. Random-effects models were computed to estimate the effect of overnight neopterin on the CAR. There was a significant positive effect of overnight neopterin levels on the CAR, indicating that overnight Th1 immune activation may predict the dynamic increase of cortisol in the morning, with higher immune levels leading to a stronger CAR. These results provide first evidence for the immunological awakening hypothesis and a potential role of the CAR in the maintenance of circadian immune rhythms, but given the small number of participants, findings should be interpreted as exploratory.

## Introduction

The glucocorticoid cortisol is the hormonal effector of the hypothalamus– pituitary–adrenal (HPA) axis and mediates a wide array of metabolic processes ranging from induction of mobilization of energy, increasing cerebral perfusion rates and local glucose utilization, enhancing cardiovascular output and respiration, redistributing blood flow, increasing substrate and energy delivery to the brain and muscles, to modulating immune function (McEwen and Seeman, 1993). Basal cortisol levels follow a well-described diurnal rhythm that peaks in the early morning and declines throughout the day reaching its low during the night (Oster et al., 2017). In addition, there is a steep increase of cortisol levels peaking about 30-45 minutes after awakening in the morning that has been termed the cortisol awakening response (CAR) and marks the transition from the rest phase to the active phase (Steptoe and Serwinski, 2016; Clow et al., 2010). The function of the CAR is still mostly unknown, but it has been hypothesized to be related to the activation of prospective memory representations at awakening and enabling an individual’s orientation about the self in time and space as well as the anticipation and adaptation of the upcoming day (Powell and Schlotz, 2012; Fries et al., 2009; Rohleder et al., 2007). Besides its proposed functions in cognitive and behavioral awakening, the CAR has also been hypothesized to play a role in an “immunological awakening” and the circadian rhythm of immune system activity (Clow et al., 2010, 2004). A better understanding of the CAR during healthy functioning will be key to comprehend the abnormal expression of the CAR and its utilization as a biomarker in psychiatric and epidemiological research (Law et al., 2015; Clow et al., 2010).

Immune system activity has been shown to be synchronized to the circadian rhythm and transitions between active and rest periods of the organism (Wang et al., 2022; Wyse et al., 2021; Cermakian et al., 2013; Lange et al., 2010; Haus and Smolensky, 1999), what may have evolved in response to temporally varying patterns of potential pathogen exposure that is increased during the active phase of an organism for example due to more social interaction, physical activity, and food intake (Scheiermann et al., 2013). Specifically, in humans, the numbers of circulating T-lymphocytes and B-lymphocytes have been shown to peak during the night, whereas recruitment to tissues happens preferentially during the day (Wang et al., 2022; Druzd et al., 2017; Cermakian et al., 2013; Scheiermann et al., 2013). Accordingly, the levels of cytokines and chemokines such as interferon gamma (IFN-γ), tumor necrosis factor alpha (TNF-α, neopterin, interleukin (IL)-1 beta, IL-2, IL-6 and IL-12 follow a similar circadian pattern with peaks during the night (Seizer et al., 2023; Cermakian et al., 2013; Bollinger et al., 2011; Born et al., 1997). For more detailed descriptions of these circadian immune rhythms, we point the readers forward to existing reviews, e.g., Wang et al (2022), Labrecque & Cermakian (2015), and Lange et al (2010).

The circadian rhythms in immune system activity seem (at least in part) to be modulated by endocrine influences, such as the proinflammatory Th1-supporting hormones melatonin, prolactin, and leptin that peak overnight, and anti-inflammatory hormones that peak during the day such as cortisol, which is one of the most potent endogenous regulators of immune activity (Oster et al., 2017). Besides the influence of cortisol on immune activity, pro-inflammatory cytokines like IL-1, IL-6, and TNF-α can surpass the blood-brain barrier and stimulate the HPA axis as part of a self-regulatory feedback loop (Turnbull & Rivier, 1995). Thus, as it has been shown that the CAR is mainly driven by hormonal release (ACTH) from the pituitary (Fries et al., 2009), inflammatory states might also influence the CAR. So-far studies that investigated the relationship of the CAR and immune system activity found the CAR to be negatively related to serum IL-6 sampled later at the same day of CAR assessment (Corominas-Roso et al., 2017), and plasma IL-6 sampled on a different day (DeSantis et al., 2012). Further, first waking salivary cortisol levels were negatively related to salivary IL-1β measured in the same sample (LaVoy et al., 2020), and plasma TNF-α sampled on a different day (DeSantis et al., 2012). However, all these studies determined cytokine levels either simultaneously with or following cortisol levels, which gives limited insights into possible roles of immune system activity as a modulator of the CAR.

The present study aims to provide pilot data on the influence of immune activity on the CAR. There are some features of the CAR that are relevant to consider in its investigation. For one, expression of the CAR on a given day is determined mostly by state variables and situational factors (62-82%), while stable, trait-like factors only account for 15-37% (Ross et al., 2014; Hellhammer et al., 2007). Thus, as a strategy to increase the reliability of results in CAR studies, it has been recommended to increase the number of study days per participant (Segerstrom & Boggero, 2020). Based on these considerations we applied an intensive longitudinal design adapted from the integrative single-case study design, which provides a framework for time-series research in psychoneuroimmunology (Schubert, 2025). Thereby each participant collected urine overnight (immune marker) and saliva upon awakening (CAR) daily over a period of one month. This approach has several strengths: (1) within one subject the trait characteristics (e.g., age, sex, body-mass-index) remain constant, what enables to focus the analysis on the influence of state variables (i.e., nightly immune activation) and within-subject variation; (2) the use of a repeated-measurement design reduces the impact of novelty effects; (3) multiple study days per subject increase the reliability of measurements; and (4) conducting sampling during everyday life increases ecological validity and avoids problems in lab-to-life generalizability (Schubert, 2025; Seizer et al., 2024; Löchner et al., 2024; Stalder et al., 2009).

## Methods

### Study design

The collection period lasted 30 days for each of the three healthy participants (2 female, 1 male; all in their mid-twenties). During this period the participants collected their urine overnight to determine nightly immune system activity and three saliva samples in the morning to determine the CAR (0-, 30- and 45-minutes post-awakening). All urine and saliva samples were stored immediately after collection at -20°C by the participants. After the 30-day collection period the samples were stored at -80°C until further analysis. Moreover, the participants wore a smart headband (Dreem 3) during the night to objectively determine the timepoint of awakening (Arnal et al., 2020). To avoid erroneous data from delayed sampling, all study days with a delay greater than 5 minutes between awakening and first saliva sampling will be excluded from the analysis (Stalder et al., 2022; Smyth et al., 2013). All subjects gave informed consent to their participation and to the publication of data. The Ethics Committee of the University Hospital Tübingen approved the study.

### Timepoint of awakening

For objective determination of the timepoint of awakening, a wireless headband was used (Dreem 3). This device records physiological data in real-time. Brain cortical activity is measured via five EEG dry electrodes and further, head position, movements, and breathing frequency are measured via a 3D accelerometer located over the head. The device has an embedded algorithm, which provides sleep staging for each 30s interval. This automatic sleep staging is conducted according to classification of the American Academy of Sleep Medicine criteria and yields performances like that of an expert consensus rating using medical-grade polysomnography data (Arnal et al., 2020). The participants put the headband on when they go to bed and wear it overnight until the final awakening in the morning.

### Cortisol awakening response

In the measurement of the CAR, we followed previous guidelines (Stalder et al., 2022). Saliva was sampled in the morning via passive drool three times after awakening (0, 30 and 45 minutes). The subjects were instructed to refrain from brushing their teeth, keep physical activity to a minimum and take nil-by-mouth apart from water until the morning sampling routine is completed. Cortisol levels in saliva were determined using enzyme-linked immunosorbent assays (ELISA; TECAN, RE52611) following the manufacturer’s guidelines. To determine the dynamic increase of the CAR each morning, the area under the curve with respect to increase for all three samples (AUC_I_; Pruessner, 2003) and the mean increase between the first and the following samples (MnInc; Miller et al., 2018) were calculated.

### Urinary neopterin measurements

Urine samples were collected overnight. Thereby, the participants were instructed to empty their bladder and discard the urine before going to bed and then collect their first morning urine after waking up. Urinary neopterin concentrations were determined by high-performance liquid chromatography using an Agilent 1100 system (Agilent Technologies, Santa Clara, CA, USA), using a previously described method with minor modifications (Hausen et al., 1982). Neopterin is released by human monocyte-derived macrophages and dendritic cells, mainly in response to stimulation by IFN-γ, a pro-inflammatory cytokine. Elevated neopterin levels reflect cellular immune activation, and neopterin is widely used as a biomarker for non-specific systemic inflammation (Stuart et al., 2020; Fuchs et al., 1992). Urinary neopterin levels are expressed as the overnight excretion rate (in µmol neopterin per hour), which is calculated by multiplying neopterin concentrations with the total overnight urine volume and dividing this total overnight neopterin output by the time of sampling in hours (Waikar et al., 2010).

### Data analysis

The influence of overnight neopterin levels on the CAR, as AUC_I_ and MnInc, and on the cortisol levels in the first awakening sample were estimated in random-effects regression analyses. Prior to estimation, neopterin levels were centered on person-means to focus the analysis on within-participant changes. Moreover, another model was estimated that additionally included the CAR of the previous day to control whether an elevated CAR one day may be related to neopterin levels that night, and to an elevated CAR the next day. In further exploratory analyses, we estimated the same models in an idiographic approach for each participant separately using only the data from one participant at a time.

## Results

There was high compliance for the collection of saliva samples (M = 94.67 %, SD = 7.57 %) and urine samples (M = 93.33 %, SD = 5.77 %). A total of 12 sampling days (14 %) were excluded from the analysis because the delay between objective awakening and first saliva sampling exceeded the 5 min criterion. This led to a total of 221 cortisol samples, 84 urine samples, and 75 valid days for the analyses across all participants.

There were significant positive effects of overnight neopterin levels on the CAR as AUC_I_ (b = 0.15, 95% CI = [0.01; 0.29], p = .032) and on the CAR as MnInc (b = 0.01, 95% CI = [0.00; 0.01], p = .041), indicating that higher neopterin levels may be associated with a greater increase in cortisol following awakening. In subsequent analyses, a model with the previous-day CAR as an additional predictor was estimated to control for autoregressive influences. However, the directions and significance of the effects of neopterin were unchanged by the inclusion of this autoregressive predictor. Further, idiographic models using the data from each participant separately were estimated. The magnitude of the effects varied across participants, but the directions remained the same as in the group model.

**Table 1.** Associations Between Overnight Neopterin Levels and the Cortisol Awakening Response.

| Outcome |  | **b** | **SE** | **95% CI** | **t** | **p** |
| --- | --- | --- | --- | --- | --- | --- |
| CAR as $AUC_i$ | Intercept | 61.81 | 9.92 | [42.26, 81.36] | 6.23 | 0.00 |
|  | Neopterin | 0.15 | 0.07 | [0.01, 0.29] | 2.17 | .032 |
| CAR as MnInc | Intercept | 1.97 | 0.37 | [0.00; 0.01] | 5.35 | 0.00 |
|  | Neopterin | 0.01 | 0.00 | [0.00, 0.01] | 2.05 | .041 |

*Note.* Coefficients are unstandardized estimates from random-effects regression models. Overnight urinary neopterin was person-mean centered, so estimates represent within-participant associations. Positive coefficients indicate that higher-than-usual overnight neopterin levels were associated with a greater next-morning cortisol awakening response. $AUC_i$ = area under the curve with respect to increase; MnInc = mean increase; CI = confidence interval.

## Discussion

In this pilot study, three healthy participants collected overnight urine samples to assess Th1 immune activation and saliva samples after awakening to determine the CAR over a period of one month during everyday life. This intensive longitudinal design enabled us to investigate within-person associations between nightly immune activity and next-morning cortisol dynamics while minimizing confounding by stable between-person differences. Overnight immune activation, measured by urinary neopterin levels, significantly predicted the magnitude of the CAR across participants, with higher neopterin levels being associated with a stronger increase in cortisol after awakening. These findings provide preliminary support for the immunological awakening hypothesis proposed in the CAR literature (Clow et al., 2010, 2004), suggesting that the CAR may be involved not only in cognitive and behavioral mobilization after sleep, but also in the transition from nighttime to daytime phase of circadian immune regulation.

This interpretation is biologically plausible in light of prior work showing that human immune activity exhibits pronounced circadian rhythmicity. During the night, several markers of Th1-type and pro-inflammatory immune activity increase, including IFN-γ-related pathways, neopterin, and several cytokines, while glucocorticoid activity is relatively low (Wang et al., 2022; Cermakian et al., 2013; Lange et al., 2010; Haus & Smolensky, 1999). In parallel, cell trafficking also shows marked day–night organization, with circulating lymphocyte subsets peaking during the night and tissue recruitment occurring more strongly during the active phase (Druzd et al., 2017; Dimitrov et al., 2009). Against this background, the CAR may represent one endocrine mechanism that helps terminate or recalibrate this overnight immune state at the transition to wakefulness. In this framework, higher overnight immune activation would require a stronger post-awakening glucocorticoid signal to facilitate the shift toward the daytime immune setpoint. Although this interpretation remains speculative, the present findings are consistent with such a model.

Several limitations need to be considered. First and foremost, the sample comprised only three participants. Although the repeated-measures design yielded comparatively rich time series and supports estimation of within-person associations, the findings remain preliminary and cannot be generalized to the wider population. Replication in larger samples is essential. Second, neopterin reflects only one aspect of immune activity - namely IFN-γ associated cellular immune activation - and therefore does not capture the full complexity of immune processes (Stuart et al., 2020; Fuchs et al., 1993). Other inflammatory mediators, particularly cytokines and chemokines, may show different or additional associations with the CAR. Third, urinary neopterin provides an integrated overnight signal, which is advantageous for capturing cumulative immune activation, but does not allow fine-grained conclusions about the precise timing of immunological changes across the night. Fourth, although the within-subject design reduces confounding by stable person-level characteristics, time-varying factors such as sleep quality, sleep duration, perceived stress, physical activity, menstrual cycle phase, or subclinical infection may still have influenced both neopterin and cortisol and should be assessed more comprehensively in future studies.

Finally, the present findings have implications for the interpretation of the CAR in clinical and translational research. The CAR is often used as an indicator of HPA-axis functioning in psychiatric, stress-related, and somatic disorders. However, if day-to-day variation in immune activity contributes to the magnitude of the CAR, then altered CAR patterns may partly reflect altered endocrine–immune coupling rather than isolated HPA-axis dysfunction. This may be particularly relevant in conditions characterized by disturbed circadian organization and immune dysregulation (Seizer et al., 2026; Yuan et al., 2019). A better understanding of the physiological determinants of the CAR may therefore help refine its interpretation as a biomarker in psychopathology and inflammatory disease. In this sense, the present pilot data does not only support a potential immunological function of the CAR, but also underline the importance of studying endocrine and immune rhythms as interacting components of an integrated circadian regulatory system (Schubert et al., 2025).

